# NRP1 and furin as putative mediators of SARS-CoV-2 entry into human brain cells

**DOI:** 10.1101/2022.01.19.476893

**Authors:** Ashutosh Kumar, Ravi K. Narayan, Sujeet Kumar, Vikas Pareek, Chiman Kumari, Rakesh K. Jha, Pranav Prasoon

## Abstract

COVID-19 has prominent neurological manifestations including psychiatric symptoms, indicating significant synaptic pathology. Surprisingly, existing evidence suggests negligible expression of the key SARS-CoV-2 host cell entry mediators *ACE2* and *TMPRSS2* in human brain, which complicates understanding of the pathomechanisms of the neuropsychiatric manifestations in COVID-19. Recent studies suggested that an alternative host-cell entry receptor, *NRP1*, can mediate entry of *furin* cleaved SARS-CoV-2 spike proteins into the host cells. However, the role of *NRP1* and *furin* in mediating SARS-CoV-2 entry in human brain cells has been least explored and remains a lacuna in the literature. We performed an *in silico* analysis of the transcriptomic and proteomic expressions of SARS-CoV-2 host-cell entry receptors and associated tissue proteases in human brain tissue, using the publically available databases. Based on the expression analysis, SARS-CoV-2 entry in human brain cells is likely to be mediated through *NRP1* and *furin*.

**Graphical Abstract:** 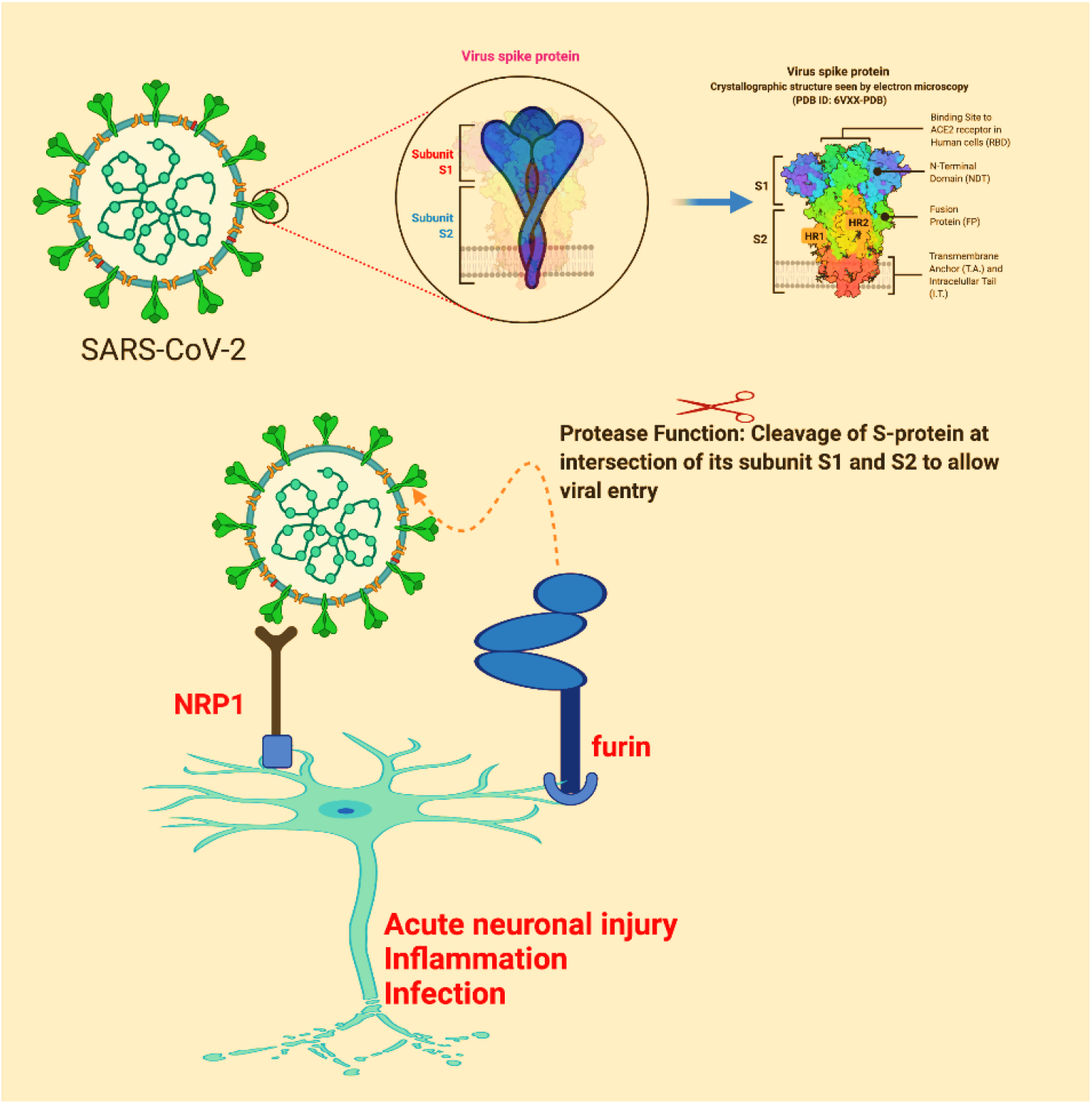

## INTRODUCTION

COVID-19 has prominent neurological manifestations as the acute and long term sequelae, including psychiatric symptoms ^1,2^. Psychiatric manifestations primarily have indicated significant synaptic pathology. SARS-CoV-2, the viral agent of COVID-19 is known to enter host-cells through a cell surface receptor, *ACE2—angiotensin converting enzyme 2* ^3^. Along with ACE2, the cellular expressions of host-proteases such as *TMPRSS2* and *furin*, which cleave the viral spike protein are considered essential for SARS-CoV-2 infection ^3^. SARS-CoV spike (S) protein uniquely contains a polybasic sequence motif, Arg-Arg-Ala-Arg (RRAR), at the S1/S2 boundary (682-685 aa). It provides a cleavage site for the host serine protease, *furin*. Inclusion of the *furin* cleavage site (FCS) is said to be an evolutionary gain in SARS-CoV-2 considering its conspicuous absence in other related SARS viruses, including SARS-CoV-1 ^3^, and has been shown to increase infectivity and tissue tropism. The emerging SARS-CoV-2 variants have accumulated mutations at the FCS potentially favoring a *furin* mediated processing ^4^. The C-terminus of the S1 protein generated by *furin* cleavage confirms to a [R/K]XX[R/K] motif, termed the “C-end rule” that can potentially bind to neuropilin receptors expressed on the host cells, and comprises of a set of host peptides known to be involved in pleiotropic physiological/pathological activities, such as axonal guidance, angiogenesis and vascular permeability, immune regulation, and survival of cancer cells. Recent studies suggested that *neuropilin-1 (NRP1)* can bind to *furin* cleaved SARS-CoV-2 spike protein and facilitate *ACE2* mediated SARS-CoV-2 entry into host cells ^5^. In addition, in higher viral titers, *NRP1* can independently mediate viral entry of into the host cells ^5,6^. Notably, blocking the *NRP1*-spike protein interaction by RNA interference or selective inhibitors reduced SARS-CoV-2 entry and infectivity in cell culture ^5,6^.

Currently, the pathomechanisms of neuropsychiatric manifestations in COVID-19 remains least understood. Among diverse contenders, a receptor based mediation of SARS-CoV-2 entry in human brain, either through olfactory or vagal nerves, or neurovascular routes remains a plausible mechanism ^7^. However, the role of *ACE2* in mediating SARS-CoV-2 entry into brain cells appears less plausible as existing evidence suggests that there is negligible *ACE2* expression in human brain tissue ^7^. There is therefore, a need to look for alternative cell-receptor, which can mediate SARS-CoV-2 entry in human brain in scarce presence of *ACE2*. Finding the key mediators of SARS-CoV-2 entry into the brain cells remains a lacuna in the literature. Multiple authors have suggested for a *NRP1* mediated mechanism behind neuropsychiatric symptoms in COVID-19 since identification of the *NRP1* as an alternative SARS-CoV-2 entry receptor in human cells (6,7). However, the studies are lacking which have studied relative distributions for the SARS-CoV-2 entry mediators and entry associated molecules in the human brain. In this study, we performed an *in silico* analysis of the publically available databases providing transcriptomic and proteomic expressions of *ACE2* and *TMPRSS2*, and *NRP1* and *furin* in human brain regions with an aim to know whether the later may have an edge in mediating COVID-19 neuropathology.

## RESULTS AND DISCUSSION

Pathogenesis of the neuropsychiatric manifestations in COVID-19 remains little understood in absence of the precise knowledge about the molecular mediators of SARS-CoV-2 entry into the brain cells. Finding the key mediators of SARS-CoV-2 entry into the brain cells may help develop target-based therapeutic strategies and suitable drug targets for treating neuro-COVID. Our study unravels relative expressions of SARS-CoV-2 host cell-entry receptors *ACE2* and *NRP1* and entry-associated host proteases *TMPRSS2* and *furin* in human brain regions providing important insights regarding the viral entry into the brain-cells and consequent neuropathology. *ACE2* and *TMPRSS2* were not-detected in transcriptomic (NX < 1) and proteomic expression analysis in any of the human brain regions, including hippocampal formation (Figs. 1a, 2).

**Figure 1.**
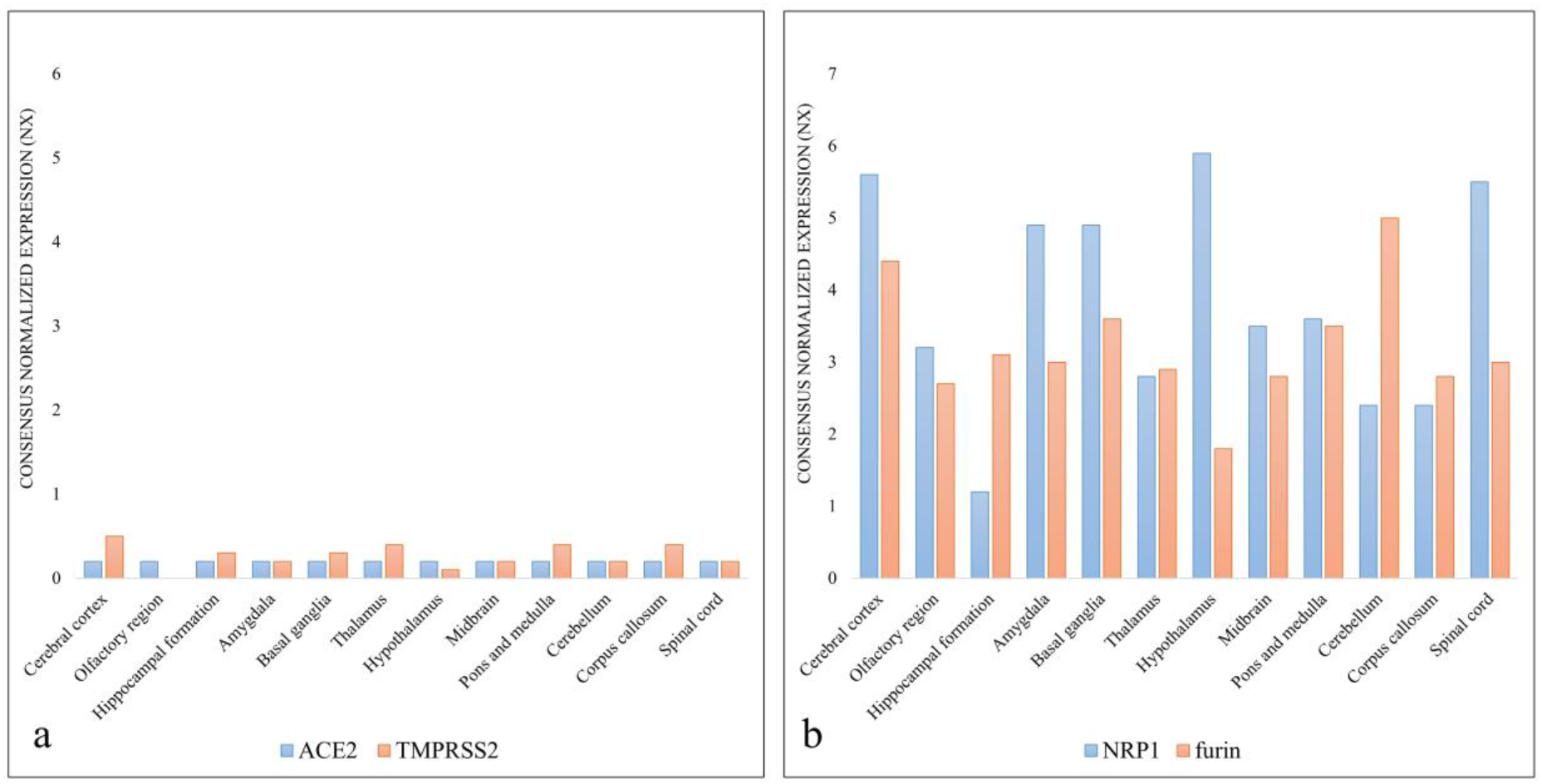
Transcriptomic expression of SARS-CoV-2 entry factors in different parts of the brain. **a**. ACE 2 and TMPRSS 2 expression in different parts of the brain; **b**. NRP1 and furin expression in different parts of the brain. **(Data source: Human Protein Atlas; https://www.proteinatlas.org/).**

**Figure 2.**
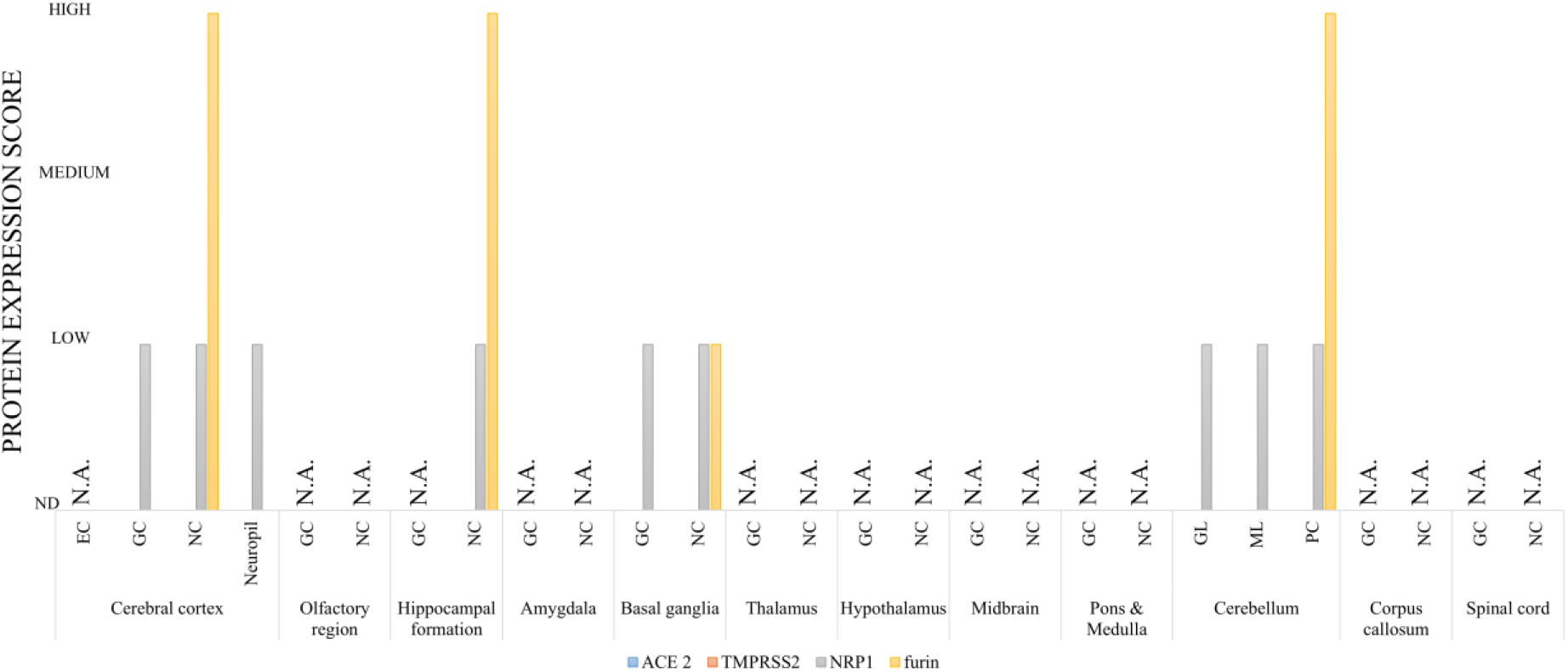
Proteomic expression of SARS-CoV-2 entry factors in different parts of the brain. (NC-Neuronal Cells; GC - Glial cells; EC - Endothelial Cells; GL-cells of Granular layer; ML - Cells of Molecular layer; PC - Purkinje cells; N.A.—Data not available). **(Data source: Human Protein Atlas; https://www.proteinatlas.org/)**.

In contrast, transcriptomic as well as proteomic expressions of *NRP1* and *furin* were detectable (NX ≥ 1) across the human brain (Figs. 1b, 2). The protein expressions of *NRP1* and *furin* were detectable in neuronal cell, neuropil, and glial cells of the brain tissue (Fig. 3).

**Figure 3.**
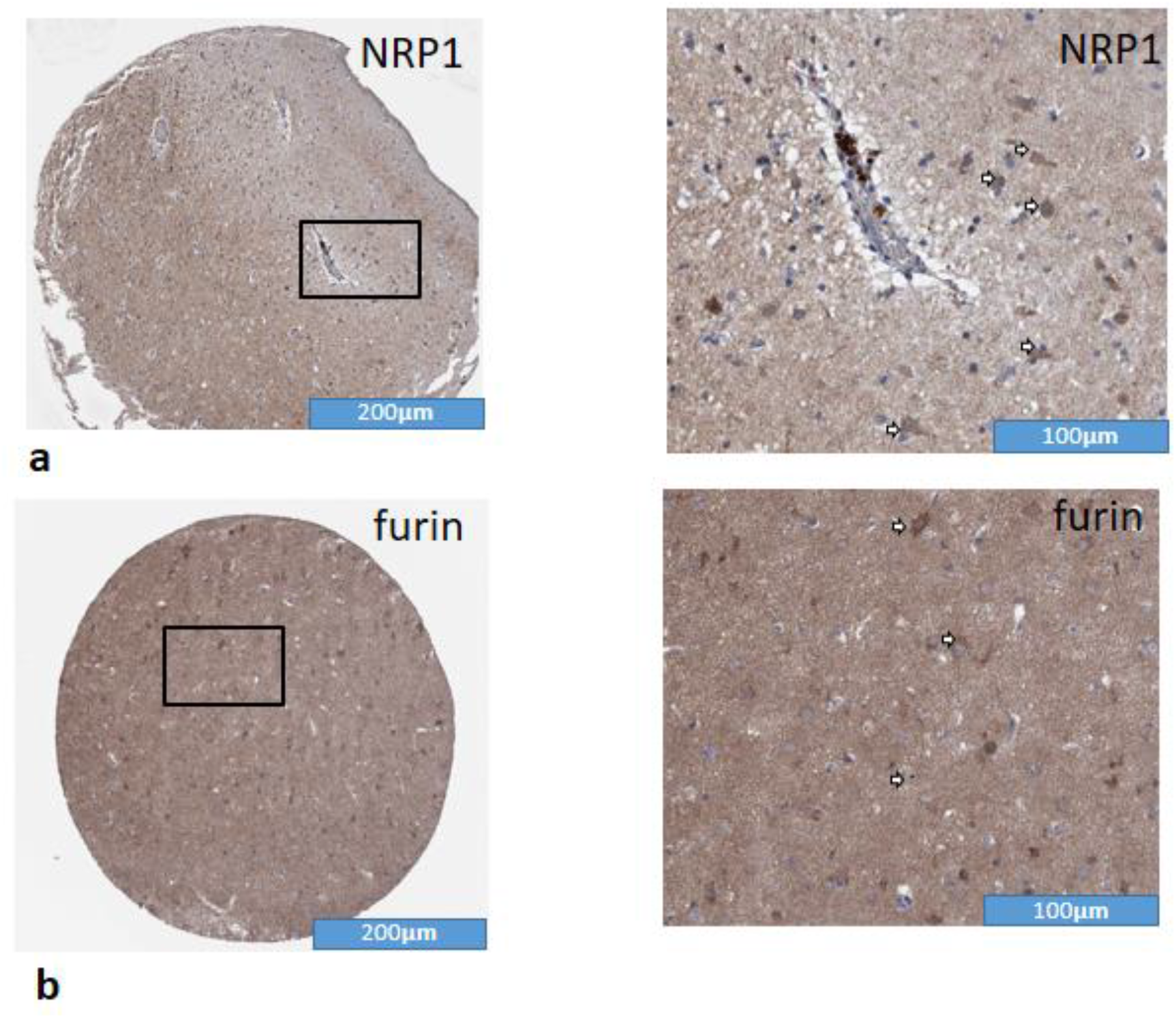
Immunohistochemical expression in the human cerebral cortex. **a.** NRP1 **b.** furin. Arrows indicate immunostaining in the cytoplasmic membrane of neuronal cells. In addition, moderate to intense staining can be also noted in the neuropil (intercellular area) and cytoplasmic membrane of glial cells (with small blue nuclei). **(Data source: Human Protein Atlas; https://www.proteinatlas.org/)**.

Based on the expression analysis of the SARS-CoV-2 entry-receptors and the entry-associated host-proteases in the human brain tissues, SARS-CoV-2 entry into human brain cells is likely to be mediated independently through *NRP1* and *furin*, instead of *ACE2* and *TMPRSS2*. The significant expression of *NRP1* and *furin* in neuronal cells, including within the olfactory epithelium and hippocampal formation, suggests that *NRP1* may have a role in the pathomechanisms of synaptic pathologies; such as, loss of the sense of smell and taste, memory loss, and multiple psychiatric manifestations in COVID-19, as the acute or long term sequelae. Notably, *NRP1* also has a known role in the regeneration and plasticity of adult neurons ^9^. It works as a receptors for *Semaphorin 3A*— an axonal guidance protein ^9^, and *vascular endothelial growth factor A (VEGF-A)* ^10^—a key molecule involved in hippocampal neurogenesis. A recent study has shown that the SARS-CoV-2 spike protein binding to *NRP1* in sensory nerves can induce analgesia by altering *VEGF-A* mediated neuronal signaling ^10^. Of note, the olfactory epithelium and the hippocampus are the known sites for the ongoing neurogenesis in the adult humans and interestingly, both of these brain regions are also implicated in COVID-19 neuropathology. A SARS-CoV-2 infection induced disruption of the ongoing olfactory and hippocampal neurogenesis and/or plasticity in COVID-19 patients remains a plausibility in view of high *NRP1* and *furin* expression in these brain regions. A disrupted neurogenesis can also explain frequent olfactory and hippocampal dysfunctions in COVID-19 cases ^11,12^. Furthermore, the SARS-CoV-2 induced neuro-immune dysfunction and neuro-inflammation can be also mediated through *NRP*, given its known role as an immune checkpoint of T cell memory and inflammation ^13^.

In conclusion, our study provides important lead for conducting *in vivo* studies considering the NRP1 and furin, as the putative mediators of SARS-CoV-2 entry into the brain-cells.

### Limitations

For certain brain regions proteomic expression data were not available to corroborate transcriptomic expressions of the studied markers. Further, the study didn’t employ any experimental evidence to corroborate the findings derived from the *in silico* analysis. The *in situ/in vivo* studies are warranted to validate our findings.

## METHODS

We performed *in silico* analysis of mRNA and protein expressions of *ACE2, NRP1, TMPRSS2*, and *furin* in human brain immune components using the tissue transcriptome and immunohistochemistry (IHC) data available in Human Brain Atlas (HBA), a sub-section of Human Protein Atlas (HPA) (https://www.proteinatlas.org/).

### External data source methods (As described by the source labs)

Estimation of mRNA expression and localization of human proteins were performed by the source laboratory using deep sequencing of RNA (RNA-seq) and IHC in normal tissue.

### IHC

The specimens containing normal-tissue were collected in accordance with approval from the local ethics committee. The specimens were derived from surgically removed tissue, normal was defined by tissue-specific morphological parameters and absence of neoplasia. Antibodies against human *ACE2* (HPA000288, CAB026174), NRP1 (HPA030278, CAB004511), *TMPRSS2* (HPA035787), and *furin* (CAB009499) were used. IHC staining was performed on normal tissue microarray using a standard protocol (https://www.proteinatlas.org/download/IHC_protocol.pdf). Protein expression score (low, medium, and high) for each protein was calculated based on the staining intensity (negative, weak, moderate or strong) and fraction of stained cells (<25%, 25-75% or >75%).

### Human brain transcriptomics

The transcriptomic data was collected from the three databases (HPA, GTEx and FANTOM5). For HPA RNAseq total RNA was extracted from the tissue samples of healthy individuals (Accession no: PRJEB4337, Ensembl: ENSG00000130234 (version 92.38) using the RNeasy Mini Kit (Qiagen, Hilden, Germany) following the manufacturer’s instructions. Experion automated electrophoresis system (Bio-Rad Laboratories, Hercules, CA, USA) with the standard-sensitivity RNA chip or an Agilent 2100 Bioanalyzer system with the RNA 6000 Nano Labchip Kit (Agilent Biotechnologies, Palo Alto, USA) was used to analyze the extracted RNA samples. Only the samples of RNA Integrity Number > 7.5 were used for the mRNA sequencing. The mRNA sequencing (read length of 2×100 bases) was performed using Illumina HiSeq2000 and 2500 machines (Illumina, San Diego, CA, USA) as per the standard Illumina RNA-seq protocol. Kallisto v0.43.1 (https://pachterlab.github.io/kallisto/about) was used for estimation of the transcript abundance. Further, the normalized Tags Per Million (TPM) for each gene from the three screened databases were calculated and each tissue was categorized for the intensity of gene expression using a cutoff value of 1 NX (as a limit for detection across all tissues). The transcriptomic expression of an analyzed protein in a particular tissue type was categorized (i) Enriched: if it had NX level at least four times higher than other tissues, (ii) Low specificity: if NX ≥ 1 in at least one tissue, (iii) Not detected: if NX < 1 in all tissues.

### Meta-data analysis

We analyzed HPA transcriptomic and proteomic data in respect of tissue specific expression of the screened proteins in human brain components. For each protein, we matched the IHC staining profile with mRNA expression data to yield an ‘annotated protein expression’ profile. Graphs were plotted and inferences were made based on the final results.

## Acknowledgements

Authors express gratitude to ‘The Human Protein Atlas (https://www.proteinatlas.org/)’ for allowing use of its metadata for this study.

## Ethics statement

An ethical clearance from the institute ethics committee for conducting the study was precluded as the open access published databases were used as the source of data.

## Author (s) Contributions

AK and VP conceived the idea. AK and RKN, and PP analyzed the data and prepared figures and tables. AK wrote the first draft. SK, RKN, CK, VP, RKJ, and PP reviewed and edited for the final draft.

